# Interrogating RNA-small molecule interactions with structure probing and AI augmented-molecular simulations

**DOI:** 10.1101/2021.09.28.462207

**Authors:** Yihang Wang, Shaifaly Parmar, John S. Schneekloth, Pratyush Tiwary

**Affiliations:** Biophysics Program and Institute for Physical Science and Technology, University of Maryland, College Park, MD 20742, USA; Chemical Biology Laboratory, Center for Cancer Research, National Cancer Institute, Frederick, MD 21702, USA; Department of Chemistry and Biochemistry and Institute for Physical Science and Technology, University of Maryland, College Park 20742, USA

## Abstract

While there is increasing interest in the study of RNA as a therapeutic target, efforts to understand RNA-ligand recognition at the molecular level lag far behind our understanding of protein-ligand recognition. This problem is complicated due to the more than ten orders of magnitude in timescales involved in RNA dynamics and ligand binding events, making it not straightforward to design experiments or simulations. Here we make use of artificial intelligence (AI)-augmented molecular dynamics simulations to directly observe ligand dissociation for cognate and synthetic ligands from a riboswitch system. The site-specific flexibility profiles from our simulations are compared with *in vitro* measurements of flexibility using Selective 2’ Hydroxyl Acylation analyzed by Primer Extension and Mutational Profiling (SHAPE-MaP). Our simulations reproduce known relative binding affinities for the cognate and synthetic ligands, and pinpoint how both ligands make use of different aspects of riboswitch flexibility. On the basis of our dissociation trajectories, we also make and validate predictions of pairs of mutations for both the ligand systems that would show differing binding affinities. These mutations are distal to the binding site and could not have been predicted solely on the basis of structure. The methodology demonstrated here shows how molecular dynamics simulations with all-atom force-fields have now come of age in making predictions that complement existing experimental techniques and illuminate aspects of systems otherwise not trivial to understand.

## 1. INTRODUCTION

RNA regulates diverse cellular processes, far beyond coding for protein sequence or facilitating protein biogenesis.^1^ In this role, RNA can interact with protein^2^ or small molecule factors^3,4^ to directly or indirectly impact gene expression and cellular homeostasis.^5^ For example, riboswitches are sequences found primarily in bacterial mRNAs that contain aptamers for small molecules and regulate transcription or translation.^6^ In humans, mutations in noncoding RNAs can directly impact neurodegenerative diseases and cancer.^7,8^ The regulatory role of bacterial riboswitches, coupled with the broader link between RNA and human disease has also led to interest in RNA as a therapeutic target for small molecules.^9–11^ However, in comparison to proteins, methods to computationally and experimentally interrogate RNA structure are relatively less mature.^12,13^ Thus, more robust tools are needed to better understand RNA structure, dynamics, and recognition of small molecules. In spite of its clear relevance to fundamental science and drug discovery efforts, the highly dynamic nature of RNA makes it difficult to study due to the diverse conformational ensembles it adopts.^14–16^ Luckily, riboswitches represent a valuable model for RNA-small molecule recognition. When ligands bind to riboswitch aptamers, the RNA undergoes a conformational change that modulates gene expression.^17,18^ The cognate ligand for a riboswitch is typically associated with its upstream gene, often part of the ligand’s biosynthetic pathway.^18,19^ This feedback mechanism, paired with well-established structural analyses, make riboswitches excellent models for RNA-ligand binding since the target ligand is known and often highly specific. ^18^

Although we can use riboswitches to model RNA-small molecule interactions, visualizing riboswitch-ligand dissociation pathways with high spatiotemporal resolution remains a challenge for in *vitro* and in *silico* experiments. Such a femtosecond and all-atom resolution is hard to directly achieve in *in vitro* experiments, while the associated timescales are several orders of magnitude too slow for molecular dynamics (MD) simulations performed even on the most powerful supercomputers. Previously, Ref. 20 used G*ō*-model simulations of PreQ_1_ riboswitches, specifically transcriptional *Bacillus subtilis* (*Bs*) and translational *Thermoanaerobacter tengcongensis* (*Tt*) aptamers. Their work suggests that in respective cases the ligand binds at late and early stages of riboswitch folding, indicating that perhaps these riboswitches respectively fold *via* mechanisms of conformational selection and induced fit. Since Ref. 20, RNA force-fields have become more detailed and accurate, but the timescales involved in RNA-small molecular dissociation continue to stay beyond reach.^21^

In order to better understand the ligand-riboswitch dissociation process, in this work we employ a combination of (i) enhanced sampling methods combining statistical physics with artificial intelligence (AI)^22–25^ and (ii) experimental techniques to measure RNA flexibility and ligand binding.^26–28^ The computational sampling techniques allow us to study the dissociation process in an accelerated but controlled manner, while the Seletive 2’ Hydroxyl Acylation analyzed by Primer Extension and Mutational Profiling (SHAPE-MaP), microscale thermophoresis (MST) and fluorescence intensity assay (FIA) provide a rigorous validation of the computational findings. Specifically, we study the *Tt*-PreQ_1_ riboswitch aptamer interacting with its cognate ligand PreQ_1_ (7-aminomethyl-7-deazaguanine) and a synthetic ligand (2-[(dibenzo[b,d]furan-2-yl)oxy]-N,N-dimethylethan-1-amine).^29^ See Fig. 1 for an illustration of the riboswitch and chemical structures of the two ligands. First, we perform 2 *μ*s long unbiased MD simulations of the ligand-bound and ligand-free systems to quantify the flexibility of individual nucleotides. Comparing the flexibility profiles from unbiased MD with SHAPE-MaP reveals some broad overall agreement, however a more detailed quantitative analysis reveals the limitations of both approaches. To simulate the dissociation process, which is too slow for MD simulations,^21^ a key challenge in most enhanced sampling methods is the need for *a priori* estimate of the dominant dissociation mechanisms as captured through the reaction coordinate (RC). However, it is very challenging to estimate such an RC without already having simulated the dissociation process, especially for flexible biomolecules like RNA.^30^ Here, we use our AI-based sampling method RAVE to directly observe the dissociation of both of these ligands from the wild-type (WT) Tt-PreQ_1_ riboswitch. RAVE automates the learning of RCs through a scheme that iterates between MD to generate data and deep learning to construct an approximate RC from this data. The MD-deep learning iteration continues until we see no further improvement in accelerating the dissociation events as measured through typical MD time to observe dissociation. The RC itself is learned as the most informative low-dimensional representation expressed as a past-future information bottleneck.^31^

**FIG. 1:**
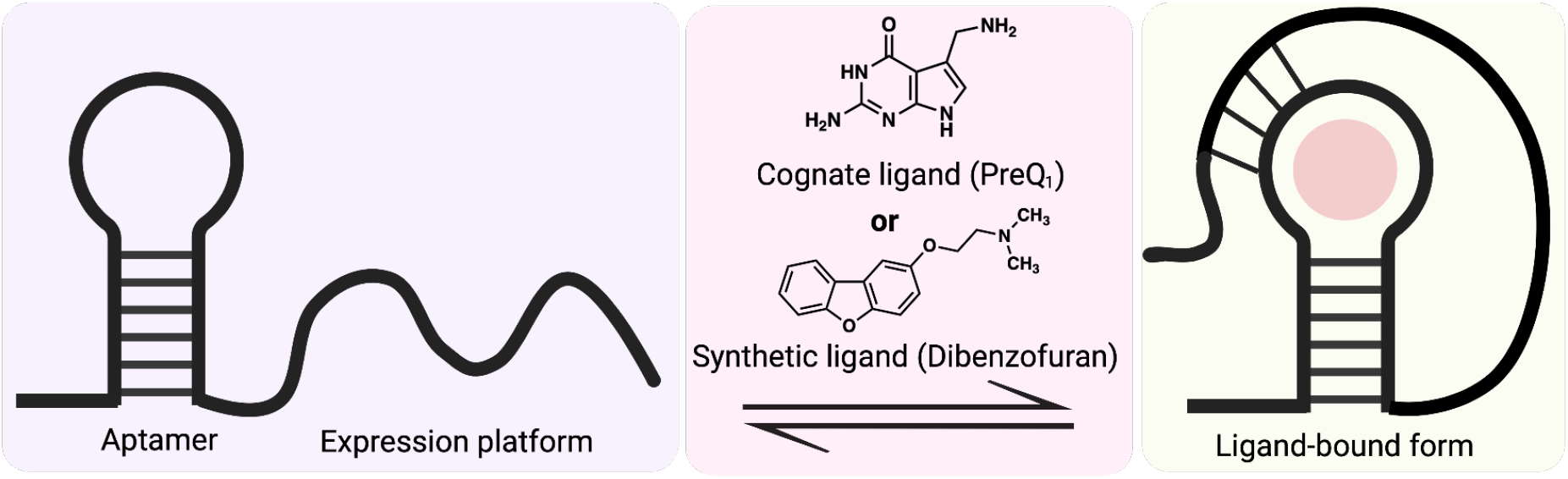
Conformational change of the PreQ_1_ riboswitch and chemical structures of PreQ_1_ and synthetic dibenzofuran ligands.

By efficiently sampling multiple independent riboswitch-ligand dissociation events for both cognate and synthetic ligands, we are able to reproduce relative binding affinity measurements from FIA and give a qualitatively reasonable fit to flexibility measurements from SHAPE-MaP experiments, though the limitations are also strikingly obvious. Furthermore, we are able to pin-point which nucleotides play the most critical roles in dissociation. These are found to be the nucleotides U22 and A32 for the cognate and synthetic ligands respectively. On the basis of these we design and perform a set of *in vitro* mutagenesis experiments, which validate our mutation predictions. We note that these predicted mutations are unlikely to have been found simply from a structural perspective, as they are distal from the binding site and do not directly interact with either of the ligands. Our findings thus reveal the complex nature of interaction between RNA and small molecules, and might help guide the design of RNA-binding small molecules in other systems in the future. Thus, this methodology will be broadly useful in understanding increasingly complex, biologically important RNA-small molecule interactions.

## 2. RESULTS

### A. Ligand binding induces flexibility change in riboswitch aptamer

Our first test is to compare the nucleotide-specific flexibility from all-atom simulations to those from SHAPEMaP measurements. We perform 2 *μ*s unbiased MD simulations (4 independent sets of 500 *ns* each for both ligands) each for PreQ_1_ riboswitch bound with the cognate and synthetic ligand respectively. See Supplementary materials (SM) for details of simulation set-up. While these simulations are not long enough to observe ligand dissociation, they can still give a robust measure of how the flexibility varies between the two different ligand-bound systems and provide nucleotide-level insight into how specific ligand binding induces different conformation changes in the riboswitch. Specifically, we compare the fluctuations of the distances between consecutive C2 atoms obtained from these simulations with SHAPE-MaP measurements (see SM for details of SHAPE-Map). We performed SHAPE-MaP analyses using a PreQ_1_ riboswitch construct consisting of the entire riboswitch (both the aptamer domain and expression platform). Probing was performed using a recently reported SHAPE reagent, 2A3, that provides improved mutagenesis rates during reverse transcription. ^32^ We analyzed data in the absence of any ligand, and in the presence of either cognate ligand (PreQ_1_) or synthetic ligand. Our results were in good agreement with previous studies reporting SHAPE-MaP on PreQ_1_ riboswitches using different SHAPE reagents and a slightly shorter construct.^33^ As discussed previously,^34^ base pairing interactions are the most important determinant of RNA structural constraints, and thus base-base distance fluctuations are a crucial metric of RNA backbone flexibility. Indeed, as shown in Fig. 2, we obtain good agreement between C2-C2 distances from unbiased MD and experimentally measured SHAPE flexibilities for all 3 systems: ligand-free PreQ_1_ aptamer, cognate-ligand-bound PreQ_1_ aptamer and synthetic-ligand-bound aptamer. We would like to note that though unbiased MD simulations capture most of the fluctuations measured by SHAPE, some of the peaks showed by SHAPE are not observed in MD, which might be due to the time scale limitations of unbiased simulations or the kinetic regime measured by SHAPE reactivity. In addition to the agreement between MD and SHAPE, Fig. 2 also shows that, for all three systems, nucleotides in the ranges A13-C15, U21-A24, and A32-G33 are more flexible than other regions of the riboswitch.^33^ The mixed agreement between MD and SHAPE approaches gives some partial amount of confidence into how accurately classical force-fields^35^ accurately model the behavior of complex RNAs, and highlights the need for continued method development from both the computational and experimental sides that can facilitate more direct comparisons of nucleotide-dependent flexibility. The lack of unequivocal quantitative agreement could be due to many reasons. Firstly, while SHAPE is a measurement of flexibility, it is strictly speaking a surrogate measurement relating to the kinetics of acylation of each nucleotide. Although flexibility of each nucleotide is a main factor in determining the probability of a nucleotide being acylated, there are other factors that can determine how the SHAPE profile looks like. Secondly, our unbiased simulations are not long enough to reduce the statistical error. Finally, as pointed out by the reviewer, we only simulated the aptamer domain of the PreQ_1_ riboswitch while in experiment the whole SHAPE reactivity of the whole PreQ_1_ riboswitch was measured. The presence of other domains may also contribute to flexibility changes in the system.

**FIG. 2:**
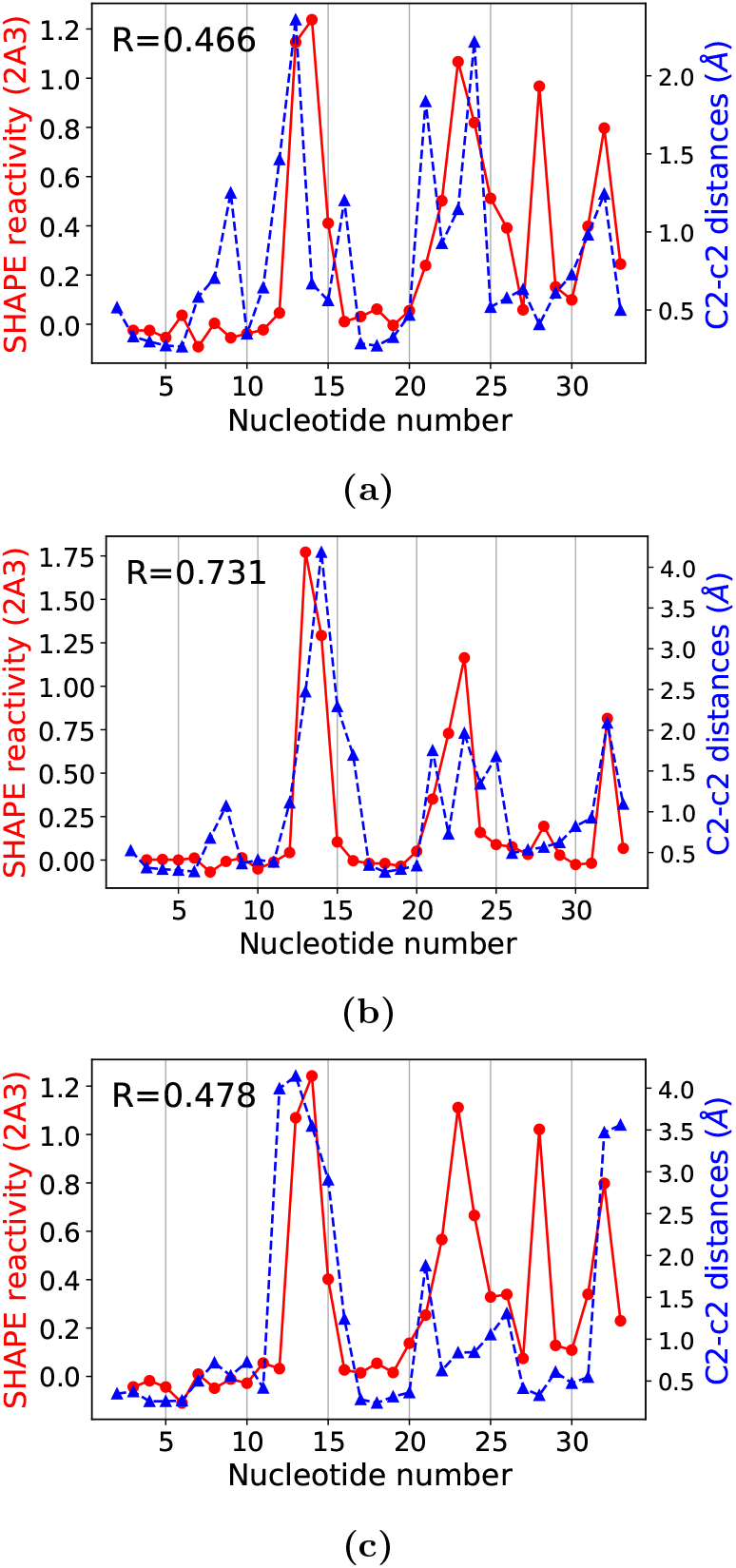
SHAPE reactivities for the *Tt* PreQ_1_ riboswitch aptamer obtained using the 2A3 reagent (red circles joined by solid lines) are compared with the fluctuations of C2 — C2 distances (blue triangles joined by dashed lines) for the PreQ_1_ riboswitch (a) in the absence of ligand, (b) bound to cognate ligand, and (c) bound to synthetic ligand. Pearson correlation coefficients *R* are shown in upper left corners.

### B. Reaction coordinates and approximate free energy profile

Encouraged by the agreement between flexibility profiles from unbiased MD and SHAPE, we next aim to calculate the free energy profiles and reaction coordinates for ligand dissociation from PreQ_1_. As these timescales are far beyond unbiased MD, here we use our recent AI-augmented MD method RAVE which learns the dominant slow degrees of freedom on-the-fly by iterating between rounds of sampling and AI. See SM for further details of the protocol. After four rounds of such iterations, we obtain converged RCs for both systems, expressed as a linear combination of different riboswitch-ligand heavy atom contacts. Fig. 3 shows different contacts and their weights as obtained from AMINO^23^ and RAVE simulations. It is interesting to note that the RC for the cognate ligand has many more components than the RC for synthetic ligand, highlighting differences between the molecules. Most of these contacts in the RC are present also in the crystal structure (PDB 6E1W, 6E1U), except carbon 5–U2, carbon 5–A23 for the cognate ligand and carbon 1–A23 for the synthetic ligand (ligand atom numbering corresponds to numbering in PDB files). We would like to highlight here that AMINO provides a basis set for RAVE to build the reaction coordinate as a function thereof, and the precise output of AMINO could differ depending on which precise trajectory was used with it. However, as shown previously in Ref. ^22^ we do not expect the final result from RAVE to significantly vary with the precise basis set fed into RAVE.

**FIG. 3:**
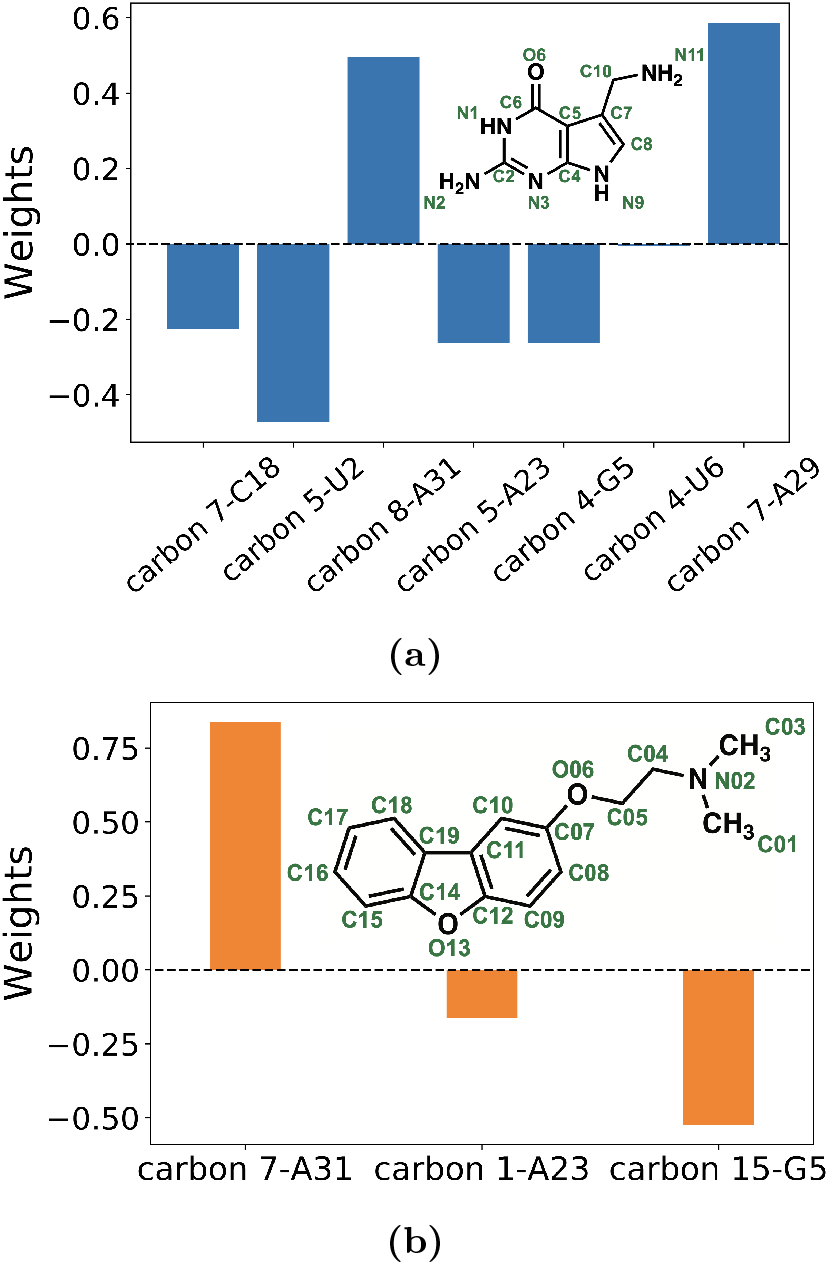
Weights for different order parameters (OPs) learned by RAVE in the corresponding RC for riboswitch aptamer with (a) cognate ligand and (b) synthetic ligand. The OPs are named in the format “ligand atom-residue name”. Ligand structures and atom names are shown in insets.

We then used this optimized RC and ran 10 independent biased MD simulations each using well-tempered metadynamics^25^ to sample the free energy profiles of the systems. The probability distribution was estimated by using the reweighting technique introduced in Ref. 36. In Fig. 4 we provide the free energy profiles as functions of the ligand-riboswitch coordination number for both systems. Note that for both systems there were 2 predominant pathways as indicated through arrows in Fig. 5. A key difference between the two paths is that in the first path, the ligand unbinds through the space in between stem 2 and the backbone connecting stem 1 and loop 1 (Fig. 5, blue arrows). In the second path, the ligand unbinds through the space in between loop 2 and stem 2 (Fig. 5, red arrows). In order to compare the binding free energies of the two systems, we combine the information from both the pathways by plotting the free energy as a function of the ligand-riboswitch shared coordination number *C*. During the dissociation process, the coordination number gradually decreases to 0 irrespective of which path is adopted during dissociation. We define the coordination number *C* as:

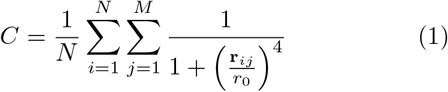

where N and M are the number of ligand and riboswitch atoms respectively. *r_ij_* is the distance between ligand atom *i* and riboswitch atom *j* and *r*_0_ is set to be 3 Å. It can be seen from Fig. 4 that when bound to the riboswitch, the cognate ligand has higher *C* relative to when the synthetic ligand is bound to it, illustrating that the synthetic ligand makes overall fewer contacts per ligand atom with the riboswitch, as also observed in crystal structures (PDB 6E1W and 6E1U). Though we were not able to simulate the rebinding of ligands to get a converged free energy profile, the probability distributions used to construct these free energies are obtained by averaging over multiple independent dissociation events. This leads to the relatively small error bars shown in Fig. 4, which indicate that we can use the free energy profile to give a rough estimation of the strength of binding affinity. By considering *C* ≲ 5.6 as the unbound states, we can calculate that the binding affinity difference between bound and unbound state is 2~4 kcal/mol higher for the cognate ligand than that for the synthetic ligand. We however caution the reader that this is an approximate free energy profile in absence of strict rebinding events during metadynamics. In principle, specialized methods^37,38^ can be used to address this limitation and will be the subject of careful investigation in future work. This is in quantitative agreement with relative affinity measurements (3.0 ± 1.4 kcal/mol) through fluorescence titrations as reported in Ref. 29.

**FIG. 4:**
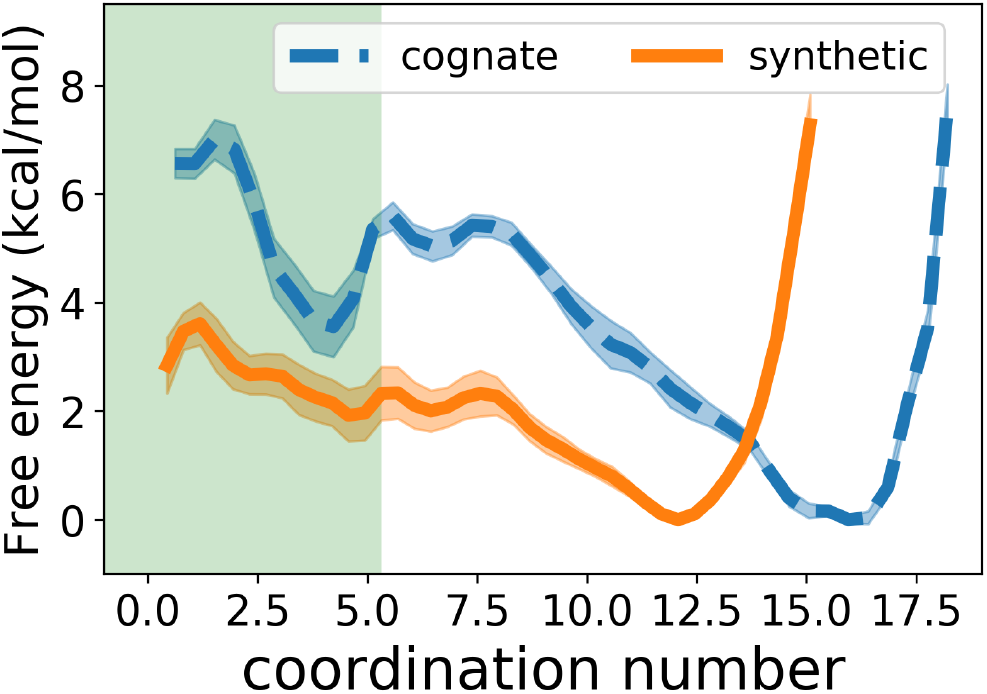
Free energy profiles of riboswitch with different ligands, Coordination number 0 (see Eq. 1) indicates ligand fully dissociated from riboswitch. The error bars are shown by the highlighted colors. The approximate range of coordination number corresponding to the unbound pose is highlighted in light green color (*C* ≲ 5.6).

**FIG. 5:**
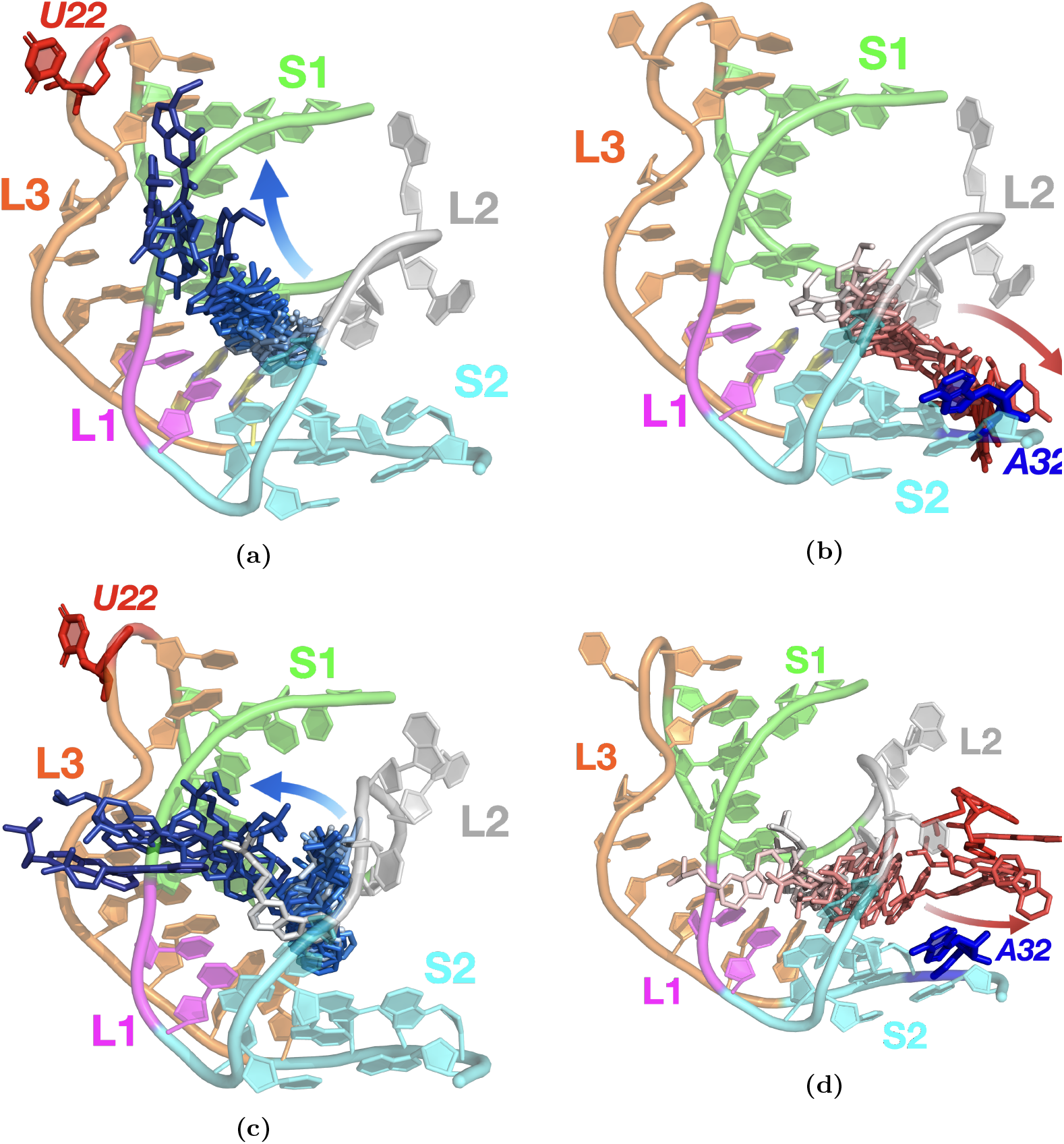
Dissociation pathways for (a)-(b) cognate ligand and (c)-(d) synthetic ligand. Stem 1 (S1), stem 2 (S2), loop 1 (L1), loop2 (L2), and loop (L3) are colored green, cyan, magenta, gray, and orange respectively. During the dissociation processes, the ligands at different time frames are colored from white to dark blue and white to red, for the top pathway and bottom pathway respectively. The dissociation pathways are indicated with arrows. The predicted mutation sites U22 and A32 are highlighted in red and blue respectively.

### C. Predicting critical nucleotides for mutagenesis experiments

We now analyze our 10 independent dissociation trajectories each obtained for the synthetic ligand bound and cognate ligand bound systems to predict critical nucleotides in the riboswitch that play a role in the dissociation process. We are interested in not just nucleotides we could have predicted from looking at the static structures of ligand-bound complexes, but also in nucleotides distal from the binding site. Those nucleotides might play an essential role in determining the structural ensemble of a riboswitch. Thus, changing the properties of those nucleotides can alter the interaction between ligand and other nucleotides. Such critical nucleotides are hard to predict from a purely structural perspective, but our all-atom resolution dissociation trajectories are ideally poised to make such predictions. Predictions made in this section are then validated through mutation experiments reported in Sec. 2D.

From our collection of dissociation trajectories, we observe that dissociation mainly happens through two pathways (Fig. 5) and that for the cognate ligand, 7 out of 10 simulations went through the top/blue pathway, while for the synthetic ligand, 6 out of 10 simulations went through the bottom/red pathway. These observations indicate how two ligands interact with the riboswitch differently before fully dissociating from the pocket. To more rigorously quantify the role played by different nucleotides, we monitor the change in the RMSD of every nucleotide in the enhanced MD simulation. In principle, a comprehensive study of ligand-receptor interactions should also include simulations of the rebinding events. As previously stated, that will require the implementation of specialized methods and thus is not explored in this study. Nonetheless, dissociation trajectories should be able to give a good sampling for bound state ensembles, allowing us to make predictions about the effects of mutations on binding affinity. Our motivation here is that the nucleotides showing greatest relative movement during the dissociation process are the ones most likely to be impacted once mutated. As shown in Supplementary Fig. 2, every nucleotide’s RMSD changes with the coordination number *C* which captures the extent of dissociation. These changes indicate different conformations the riboswitch adopts during ligand dissociation. Through this procedure we identify U22 and A32, highlighted in Fig. 5, to be two such critical nucleotides for the cognate and synthetic ligands respectively (see SM for further details of this protocol). Fig. 6 shows the changes in the RMSDs of these two selected nucleotides with different coordination numbers calculated from the enhanced MD trajectories. For U22, in RAVE simulations the RMSD increases much more for the cognate ligand-bound system, than for the synthetic ligand-bound system. However, the reverse is found to be true for the nucleotide A32, which displays more enhanced movement during the dissociation of the synthetic ligand bound complex relative to the cognate ligand bound complex.

**FIG. 6:**
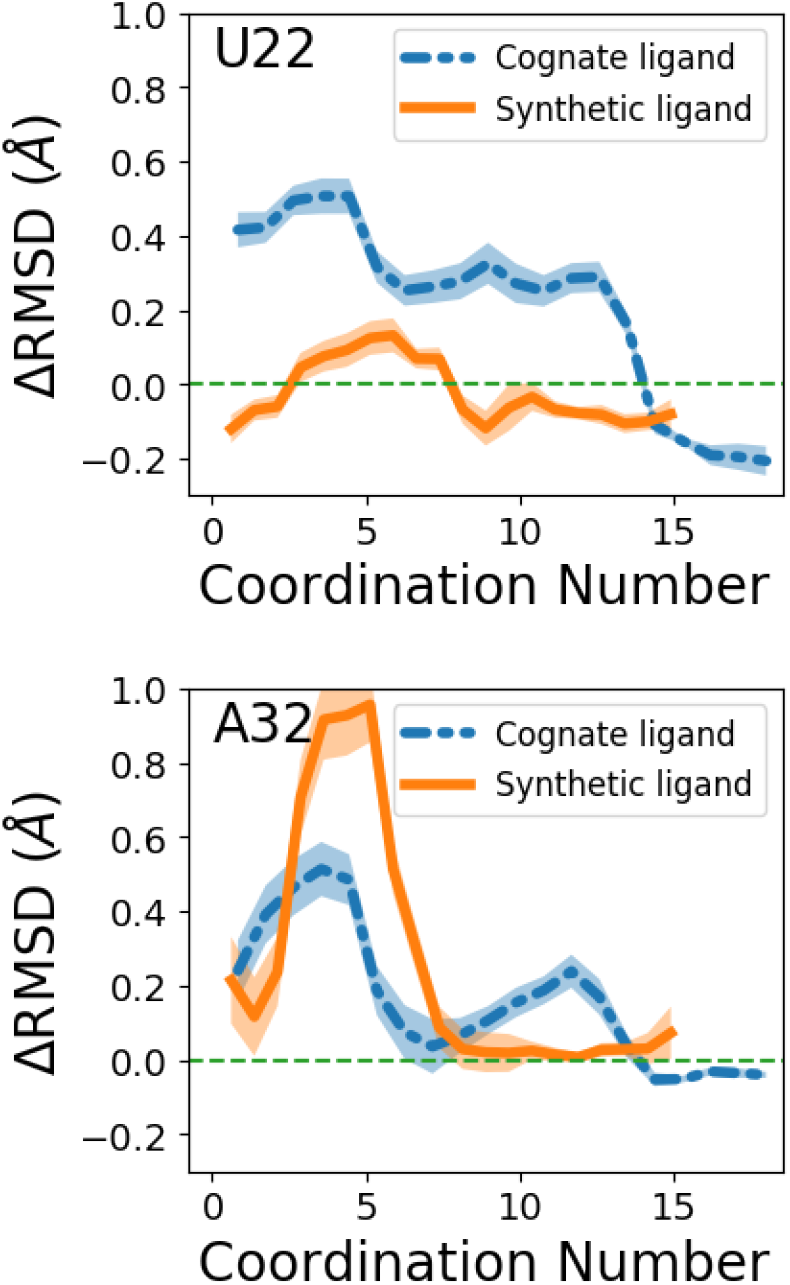
Critical nucleotides prediction. Change in RMSD for nucleotides U22 and A32 during the dissociation process, relative to RMSD variation in the apo state. RMSD profiles as function of coordination number are shown by blue dashed line and orange solid line for systems with cognate ligand and synthetic ligand separately. The change in RMSDs is calculated by subtracting the mean RMSDs of the corresponding nucleotide in apo system from the RMSDs measured in the biased simulations.

On the basis of the above observations, we can predict that for the cognate ligand, mutating U22 should display more tangible effects on the strength of the complex than mutating A32. For the synthetic ligand, we predict an opposite trend – mutating A32 should have a more pronounced effect on the interaction relative to mutating U22.

### D. Validating predicted critical nucleotides through mutagenesis experiments

In Sec. 2 C we predicted on the behalf of our dissociation trajectories that mutating nucleotides U22 and A32 will have differing effects on the cognate ligand-bound and synthetic ligand-bound systems. Also, we would like to note that, as we can see from Fig. 6, the RSMD changes appear in the range of coordination number that corresponds to the bound state. These changes imply the role of these nucleotides in determining the bound state interactions and suggest that the effects of mutations can be reflected by changes in binding affinity. Here we report *in vitro* tests of our predictions. As shown in Fig.7, we measure the equilibrium dissociation constant *K_D_* for six different complexes, namely the two wild-type complexes, U22A-cognate ligand, U22A-synthetic ligand, A32U-cognate ligand, and A32U-synthetic ligand. We also performed *in silico* mutagenesis experiments to validate that the mutations changes the bound state ensemble of both systems.

**FIG. 7:**
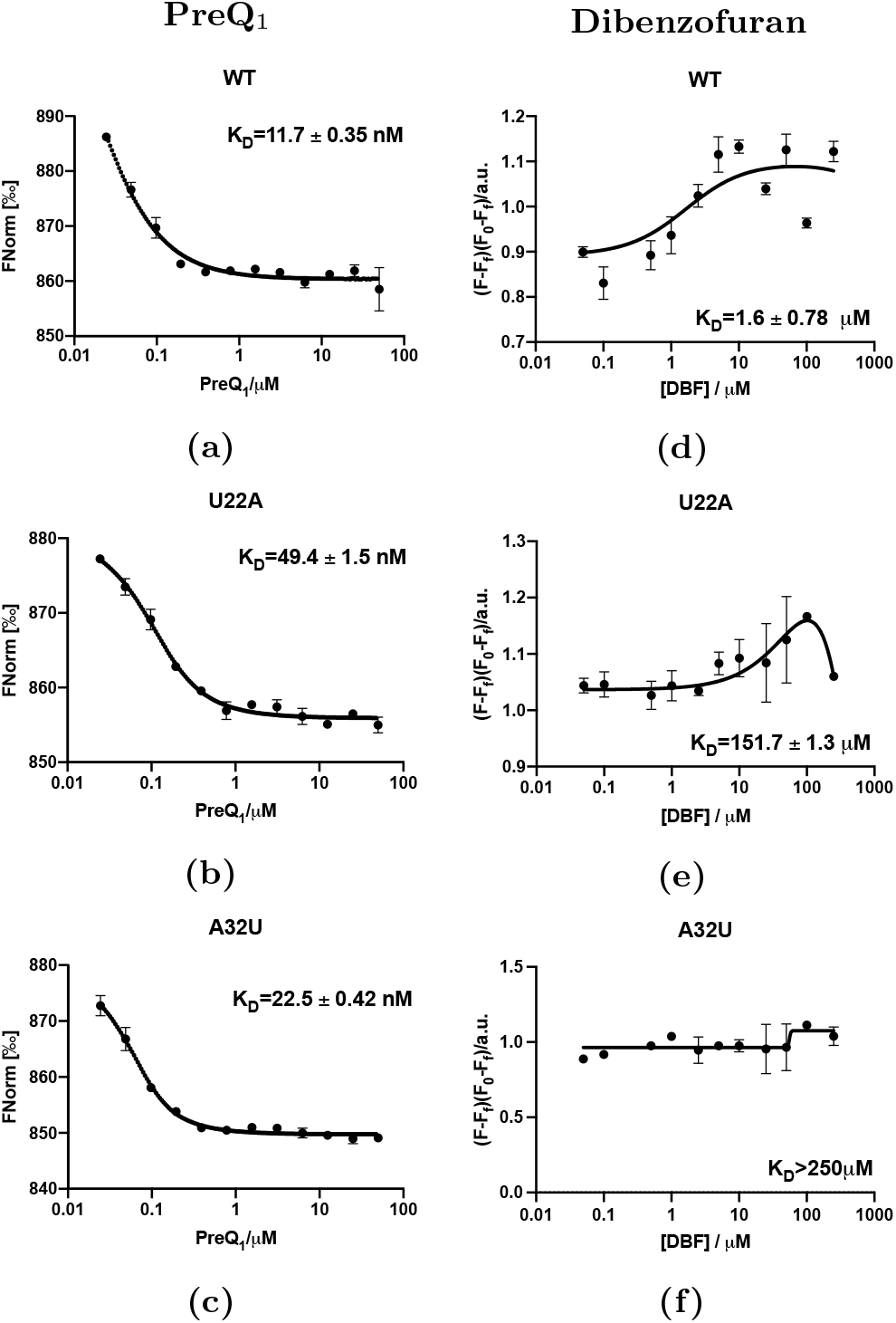
Affinity measurement of PreQ_1_ (cognate) and dibenzofuran (synthetic) ligands binding to WT *Tte* riboswitch and U22A and A32U mutants. MST of 5’-cy5-labelled (a) WT *Tte*, (b) U22A, and (c) A32U mutants in the presence of PreQ_1_ ligand. FIA of 5’-cy5-labelled (d) WT *Tte*, (e) U22A, and (f) A32U mutants in the presence of dibenzofuran. Error bars represent the standard deviation of three replicate experiments.

To experimentally test whether the A32U and U22A mutations impacted *K*D values, we used microscale thermophoresis (MST). *K_D_* values were measured using fluorescently labeled wild type, A32U, and U22A PreQ_1_ riboswitch aptamer constructs for the cognate ligand. For PreQ_1_, we observed a *K_D_* of 11.7 ± 0.35 nM for the WT construct. In contrast, the A32U mutant had a *K_D_* of 22.5 ± 0.42 nM while the U22A mutant had a *K_D_* of 49.4 ± 1.5 nM. For the synthetic ligand, changes were observed in fluorescence of the labeled RNA upon incubation with the ligand, and thus a fluorescence intensity assay (FIA) was used to measure *K_D_* values rather than MST. Using FIA we observed a *K_D_* of 1.6 ± 0.7 *μ*M for the wild type aptamer. In contrast, both the U22A and A32U mutant constructs displayed significantly weaker *K_D_* values (150 ± 1.3 *μ*M, and ≥ 100 *μ*M respectively). These *K_D_* measurements directly validate the findings from Sec. 2 C. There we had predicted that mutating U22 will effect the cognate ligand-bound system more than mutating A32, and indeed we find that *K_D_* for U22A is 5 times weaker than WT, while only 2 times weaker for A32U. At the same time, we had predicted that mutating A32 will effect the synthetic ligand-bound system more than mutating U22, and indeed we find that while the *K_D_* for U22A is 75 times weaker than WT, there is effectively no tangible association at all for A32U with the synthetic ligand.

The *in silico* experiments also support that the mutations alter the bound state ensemble and give more insights into the interaction change in the bound state after the mutations. As shown in Supplementary Fig. 3, the free energy profiles of the bound state ensemble of the mutated system differ from that of the wild type. Further analyses of the hydrogen bond interactions between ligand and riboswitch show that even though nucleotides 22 and 32 do not directly form hydrogen bonds with both ligands in the bound pose as seen in the crystal structure, interactions between ligands and other nucleotides are still altered when they are mutated, and can be seen in simulating the bound pose ensemble of states. See the Supplementary Material for detailed discussions of these simulations.

## 3. DISCUSSION

Over recent years a consensus view is emerging that a superior understanding of biomolecular structural dynamics, and not just static structures, can be fruitful in predicting key emergent biochemical and biophysical properties.^5,39,40^ However, in spite of the importance of the problem, tools to study biomolecular structural dynamics are still limited in their scope. This is true for proteins but even more so for RNAs, where it is especially challenging to determine the ensemble of possible structures that the RNA can adopt, and pinpoint the specific importance of different structures.^39^ In this work we demonstrate that this is indeed feasible - and how one can obtain detailed and robust insights into RNA structural dynamics through AI-augmented molecular simulations that maintain all-atom resolution for RNA, ions and all water molecules. All our predictions are validated through a gamut of complementary and experimental techniques that measure RNA flexibility and ligand binding. We specifically focused on the Tt-PreQ_1_ riboswitch aptamer system in its apo state and bound to two different ligands, where we first showed that 2 *μs* long unbiased MD simulations with classical all-atom force-fields for RNA and water can provide the same flexibility profile as measured in SHAPE experiments. Since the timescales for ligand dissociation are far slower than MD can typically access, we used the RAVE simulation method^22^ that enhances sampling along a low-dimensional reaction coordinate learned on-the-fly through the past-future information bottleneck framework. Through RAVE we obtain multiple independent dissociation trajectories for both cognate and synthetic liganded systems, demonstrating how the cognate and synthetic ligands invoke different aspects of the riboswitch’s flexibility in order to dissociate. On this basis, we were able to predict pairs of mutations which would be expected to show contrasting behaviors for the cognate and synthetic ligands. The ability to make such predictions accurately and efficiently is of vital importance, as mutations that can disrupt RNA structural dynamics and ligand recognition processes have been linked to numerous human diseases.^39,41^ Our predictions for the Tt-PreQ_1_ aptamer were validated by designing mutant aptamers and performing affinity measurements.

Our work thus demonstrates a pathway to gain predictive insights, as opposed to retrospective validation, from enhanced molecular dynamics RAVE simulations that maintain all-atom resolution while directly observing processes as slow as ligand dissociation. While the focus in this work was on predicting the thermodynamically averaged aspects of riboswitch flexibility, in future work it should also be possible to directly calculate rate constants using complementary approaches and for more complicated RNA systems. More broadly, this work indicates that long time scale MD simulations can accurately model ligand dissociation from RNA, and provide validatable hypotheses about structural features that impact small molecule recognition of RNA. Thus, this strategy will be widely applicable not only to study natural systems such as riboswitches, but to more completely understand the structural features and dynamics that govern how synthetic small molecules interact with RNA.

## Supporting Information

The supporting Information provides details of various methods and further analysis. These include details of:

1. *In vitro* Transcription of TTRS RNA
2. Acylation of RNA *in vitro*
3. Primer Extension and Mutational profiling (MaP) reverse transcription of RNA
4. SHAPE-MaP Library preparation, sequencing, and data analysis
5. Microscale Thermophoresis
6. Fluorescence intensity assay
7. MD simulation setup
8. Flexibility measurement from MD simulations and comparison with SHAPE data
9. AI-augmented enhanced sampling method RAVE
10. Network architecture and hyperparameter setting for enhanced sampling simulation
11. Predicting critical nucleotides for mutagenesis experiments
12. *In silico* mutagenesis experiments

## Acknowledgements

The authors thank Deepthought2, MARCC, and XSEDE (projects CHE180007P and CHE180027P) for providing computational resources used in this work. Y.W. would like to thank NCI-UMD Partnership for Integrative Cancer Research for financial support. This work was supported by the intramural program of the National Institutes of Health, National Cancer Institute, Center for Cancer Research (1 ZIA BC011585 07) (J.S.S. and S.P.) Research reported in this publication was supported by the National Institute Of General Medical Sciences of the National Institutes of Health under Award Number R35GM142719 (P.T.) The content is solely the responsibility of the authors and does not necessarily represent the official views of the National Institutes of Health. The authors thank Philip Homan for help analyzing SHAPE-MaP data and Alex Wilson for help setting up the simulations. We also thank Dr. Sergey G. Tarasov and Marzena Dyba from Biophysics resource core for help with MST experiments.

## Data availability statement

The SHAPE-MaP sequencing reads were deposited into the NLM/NCBI Sequence Read Archive (SRA) under the BioProject ID PRJNA767082.

All PLUMED input files required to perform the enhanced MD reported in this paper are available on PLUMED-NEST (www.plumednest.org), the public repository of the PLUMED consortium^42^, as plumID:22.013. The GROMACS input files and raw data to reproduce the simulation results in this paper can be found in go.umd.edu/wang_ACS_central_science_2022.

---

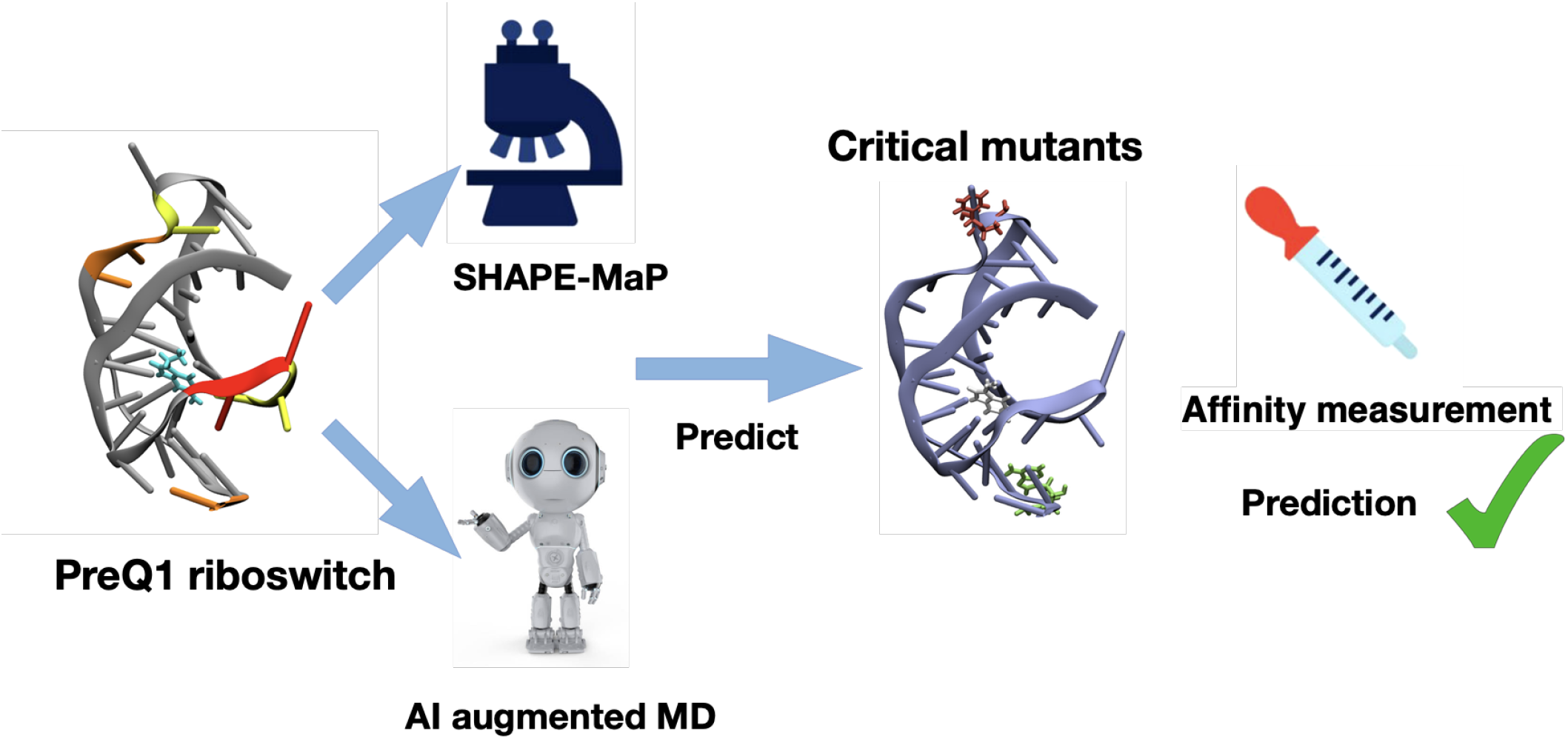
Synopsis: We use artificial intelligence-augmented molecular dynamics to study a riboswitch aptamer in complex with ligands. We obtain ligand dissociation and predict mutations that are then verified.

