## Supplementary Materials for "Interrogating RNA-small molecule interactions with structure probing and AI augmented-molecular simulations"


### 1. *IN VITRO* TRANSCRIPTION OF TTRS RNA

Template for *in vitro* transcription was generated by PCR amplification of the Ultramer® DNA Oligo (IDT) (Supplementary table 1) using Q5® High-Fidelity DNA Polymerase (NEB) following manufacturer's instructions. A full list of primers used can be found in Supplementary table 2. The RNA transcript was synthesized using the HiScribe™ T7 High Yield RNA Synthesis Kit (NEB) and purified on a denaturing (8 M urea) 10% polyacrylamide gel. The band containing the correct sized RNA was detected by UV-shadowing and excised from the gel. RNA was recovered by soaking the gel slice overnight at 4°$C$ in elution buffer (1.5 M NaOAc, 0.5 M EDTA, pH 8.0). The resulting solution was purified twice with secondary butanol and ethanol precipitated. Pure RNA was pelleted by centrifugation, washed with cold 70% ethanol, resuspended in nuclease-free water and stored at -20°$C$.

### 2. ACYLATION OF RNA *IN VITRO*

12 $\mu$l of nuclease-free water was added to 10 pmol of RNA and denatured by heating at 95°$C$ for 2 min and snap cooling on ice.[1] The denatured RNA was folded using 6 $\mu$L of 3.3× $PreQ_1$ folding buffer (165mM Tris, pH 7.5, 333mM KCl, 3.3mM $MgCl_2$) and equilibrated at 75°$C$ for 5 min and slowly cooled to room temperature for 30 min. The folded RNA was incubated with 0.5 $\mu$l each of DMSO, $PreQ_1$ (cognate ligand) and Dibenzofuran (synthetic ligand) with a final concentration of 2.5%, 10 $\mu$M and 300 $\mu$M respectively, for another 20 min. Each sample was divided into 9 $\mu$L (5 pmol each) of untreated (-) and treated (+) RNA solutions, where (-) RNA was treated with 1 $\mu$l DMSO and (+) RNA was chemically modified using 1 $\mu$l 2A3 (1 M) followed by incubation at 37°$C$ for 20 min and was quenched with 1M DTT. Completed reactions were purified using illustra G-25 columns and stored at -20°$C$ till further use.[2]

### 3. PRIMER EXTENSION AND MUTATIONAL PROFILING (MAP) REVERSE TRANSCRIPTION OF RNA

For detection of 2'-O-adducts, 9 $\mu$l of both (+) and (-) RNA were mixed, respectively, with 1 $\mu$l of dNTPs (10mM each, NEB) and 1 $\mu$l RT oligo (20 $\mu$M). The samples were incubated at 70°$C$ for 5 min and immediately transferred to ice for 1 min. To reverse transcribe the RNAs, 4 $\mu$l of first strand buffer (5×, 250mM Tris-HCl pH 8.0, 375mM KCl), 2 $\mu$l DTT (100mM), 1 $\mu$l RNase Inhibitor, 1 $\mu$l SuperScript II (SSII, Invitrogen), 1 ul MnCl2 (120 mM) were added to each tube, followed by incubation at 25°$C$ for 5 min. The mix was then incubated at 42°$C$ for 2 hours for cDNA strand synthesis and SSII was heat-inactivated by incubating at 75°$C$ for 20 min. 1 $\mu$l NaOH(10M) was added and incubated at 95°$C$ for 3 min to hydrolyze the reaction. The mix was and then transferred to ice. The cDNAs were purified using illustra G-25 columns and stored at -20°$C$ till further use.

---


* These two authors contributed equally.

### 4. SHAPE-MAP LIBRARY PREPARATION, SEQUENCING, AND DATA ANALYSIS

The libraries were prepared by the PCR amplification of the reverse transcribed cDNA (Q5® High-Fidelity DNA Polymerase, NEB). First PCR (25 $\mu$l for 10 cycles) was carried out for the amplification of the cDNA using TTRS specific forward primer (10 $\mu$M, Supplementary table 2). For unique barcoding of the libraries, another PCR was performed (50 $\mu$l for 20 cycles) using primers listed in ([1], Supplementary table 2). Each library was then purified using Zymo DNA clean and concentrator-5 (Zymo research) and eluted in 10 $\mu$l ultrapure water and were stored at -20°$C$ until sequencing. All the libraries were pooled and sequenced on an Illumina Miniseq instrument following standard sequencing protocol, outputting 2 × 150 paired-end data sets. The modification-induced mutations were analyzed using the SHAPEmapper software (https://github.com/Weeks-UNC/shapemapper2) using default parameters.[3] For the data analysis, the read depth for all the SHAPE samples (treated, untreated and denaturing controls) was required to be $> 5000$.

### 5. MICROSCALE THERMOPHORESIS

[4]100nM TTRS, U22A, A32U RNAs (IDT) were prepared in 1X $PreQ_1$ buffer (50 mM Tris, pH 7.5, 100 mM KCl, 1 mM $MgCl_2$) and annealed by heating to 75°$C$ for 5 min and then cooled to room temperature for 30 mins. A 2-fold dilution series of the cognate ligand ($PreQ_1$) was prepared using 10% DMSO in $PreQ_1$ buffer. Serial dilutions of the ligand was made with starting concentration of 50 $\mu$M. 10 $\mu$l of folded RNA was allowed to equilibrate with the ligand for 15 min at RT. Following incubation, the samples were added to premium coated capillaries and MST experiments were conducted in triplicate on a Monolith NT.115 system (NanoTemper Technologies). The results were analyzed by TJump analysis, the values obtained were normalized to the DMSO control, and plotted against the ligand concentration. The apparent dissociation constant ($K_D$) was then determined using a single-site binding model to fit the curve.

### 6. FLUORESCENCE INTENSITY ASSAY

The fluorescence intensity assay was used to determine the binding affinity of the synthetic ligand (Dibenzofuran) with wild type-TTRS and the two mutants (U22A, A32U) (Supplementary table 3). Titrations were performed using a 5'-Cy5-labeled TTRS, 5'-Cy5-labeled U22A, 5'-Cy5-labeled A32U RNA constructs purchased from IDT. The RNAs were annealed in 1× $PreQ_1$ folding buffer at 75°$C$ for 5 min and slowly cooled to RT for 30 mins. In a black 96-well plate (Costar, black side clear bottom), ligand solutions were prepared as serial dilutions at concentrations ranging from 0 to 250 $\mu$M in triplicates in 1× $PreQ_1$ buffer with a 5% final DMSO concentration and incubated at RT with gentle shaking. 5'-Cy5-labelled RNAs were added to each well to a final concentration of 100 nM in the plate. The samples were allowed to equilibrate at 37°$C$ for 30 min with gentle shaking, followed by a short spin down. The fluorescence intensity was then measured on a Synergy Mx microplate reader (BioTek) at an excitation wavelength of 649 nm and an emission wavelength of 670 nm with gain being 125. The fluorescence intensities were normalized to the values obtained for RNA only incubated with a DMSO control and were plotted against ligand concentration. The binding affinity was calculated by fitting the curve using-site total model in GraphPad Prism 8.3.1 software.[5]

| | |
|---|---|
| Ultramer® DNA Oligo sequence: | TAATACGACTCACTATAGGGCCTTCGGGCCAACTCACCTGGGT<br>CGCAGTAACCCCAGTTAACAAAACAAGGGAGGTAATTTTTCG<br>ATCCGGTTCGCCGGATCCAAATCGGGCTTCGGTCCGGTTC<br><br>(T7 promoter-5' linker-TTRS-3' linker-RT primer binding site) |

Supplementary Table 1

### 7. MD SIMULATION SETUP

The molecular replacement (MR) method has been used in the past to solve the crystal structure of the $PreQ_1$ riboswitch with a variety of ligands (PDB ID: 6e1W, 6e1U).[6] However, nucleotides 13 and 14 at L2 loop were not fully solved in the crystal structures as these positions were mutated to promote ligand binding. Using the loop structure from 3Q50[7] as a template, we filled the missing nucleotides at positions 13 and 14 using MODELLER.[8,9]

| | |
|---|---|
| **IVT:** | T7 promoter sequence underlined. |
| TT_temp_F | TAATACGACTCACTATAGGGCC |
| TT_temp_R | GAACCGGACCGAAGCC |
| **RT:** | |
| cDNA RT | GAACCGGACCGAAGCCCG |
| **SHAPE_PCRI** | |
| PCRI_F | GACTGGAGTTCAGACGTGTGCTCTTCCGATCTNNNNNCTCACCTGGGTCGCAGTAAC |
| PCRI_R | CCCTACACGACGCTCTTCCGATCTNNNNNGAACCGGACCGAAGCCCG |
| **SHAPE_PCRII** | Unique barcode sequences are underlined. |
| PCRII_FD1 | CAAGCAGAAGACGGCATACGAGATGGCCACGTGACTGGAGTTCAGAC |
| PCRII_FD2 | CAAGCAGAAGACGGCATACGAGATCGAAACGTGACTGGAGTTCAGAC |
| PCRII_FQ1 | CAAGCAGAAGACGGCATACGAGATCGTACGGTGACTGGAGTTCAGAC |
| PCRII_FQ2 | CAAGCAGAAGACGGCATACGAGATCCACTCGTGACTGGAGTTCAGAC |
| PCRII_FF1 | CAAGCAGAAGACGGCATACGAGATGCTCATGTGACTGGAGTTCAGAC |
| PCRII_FF2 | CAAGCAGAAGACGGCATACGAGATAGGAATGTGACTGGAGTTCAGAC |
| PCRII_R | AATGATACGGCGACCACCGAGATCTACACTCTTTCCCTACACGACGCTCTTCCG |

Supplementary Table 2: Primer sequences

| | |
|---|---|
| cy5-TTRS | /5Cy5/rCrUrCrArCrCrUrGrGrGrUrCrGrCrArGr<br>UrArArCrCrCrCrArGrUrUrArArCrArArArArCrArArGrGrGrArGrGrUrArArUrUrUrU |
| cy5-U22A | /5Cy5/rCrU rGrGrG rUrCrG rCrArG rUrArA<br>rCrCrC rCrArG rUrArA rArCrA rArArA rCrArA |
| cy5-A32U | /5Cy5/rCrU rGrGrG rUrCrG rCrArG rUrArA<br>rCrCrC rCrArG rUrUrA rArCrA rArArA rCrArU |

Supplementary Table 3: RNA sequences

We use the software Avogadro[10] to add hydrogen atoms to the cognate and synthetic ligands denoted HNG and HMJ respectively. ACPYPE[11] parameterizes the ligand with general amber forcefield(gaff), and ligand charges are assigned by semi-empirical method (AM1) with bond charge correction (BCC)[12], assuming a total charge of +1. NaCl is added to the system at a concentration of 0.15 mol/L, and TIP4P-D is our chosen water model.[13] For each system, its energy was first minimized with the steepest descent algorithm. Next, the system was equilibrated to 300 K over 100 ps under the NVT ensemble. The All heavy atoms of the RNA and ligand were under the positional restraints of 1000 $\mathrm{kJ \cdot mol^{-1} nm^{-2}}$ and Bussi-Parrinello velocity rescaling thermostat[14] were used to maintain the temperature. Then, the system was equilibrated under the NPT ensemble for another 100 ps with the same positional restraints. Bussi-Parrinello velocity rescaling thermostat[14] and Berendsen barostat[15] were used to separately maintain the temperature and pressure. We perform all simulations using the AMBER ff14 RNA force field[16] reparametrized by D. E. Shaw research.[17] The force field files are obtained from the GitHub repository at https://github.com/srnas/ff/tree/desres, which is maintained by "Small ribonucleic acids in silico" project hosted by the International School for Advanced Studies. Parrinello-Rahman barostat[18] and Bussi-Parrinello velocity rescaling thermostat[14] were used to simulate the NPT ensemble at temperature 300K and pressure 1 bar. Simulations were performed by using GROMACS 2016 compiled with Plumed 2.4.[19]

### 8. FLEXIBILITY MEASUREMENT FROM MD SIMULATIONS AND COMPARISON WITH SHAPE DATA

The SHAPE reactivity is known to be associated with the flexibility of the nucleotide. In our study, we followed the approaches in previous studies[20–22] to analyze the correlation between SHAPE reactivity and flexibility measurement from MD simulations. We chose two quantities that were reported to correlate with the SHAPE reactivity: the fluctuations of the distance between consecutive C2 atoms (C2-C2 distance), and the root-mean-square deviation (RMSD) for each nucleotide.

Four independent unbiased MD simulations each 500 $\mu s$ long are run starting with different randomly initialized velocities. The RMSD of each nucleotide is recorded with respect to the initial configuration, and the $i$th C2-C2 distance is recorded as the distance between C2 atoms from consecutive nucleotides $i$ and $i+1$. We calculated the Pearson correlation coefficient between flexibility measurements from MD simulations and the SHAPE-MaP measurements to quantify their correlation. Supplementary Fig. 1 shows the pairwise correlation between SHAPE-MaP measurements and the fluctuation of C2-C2 distances for different systems.

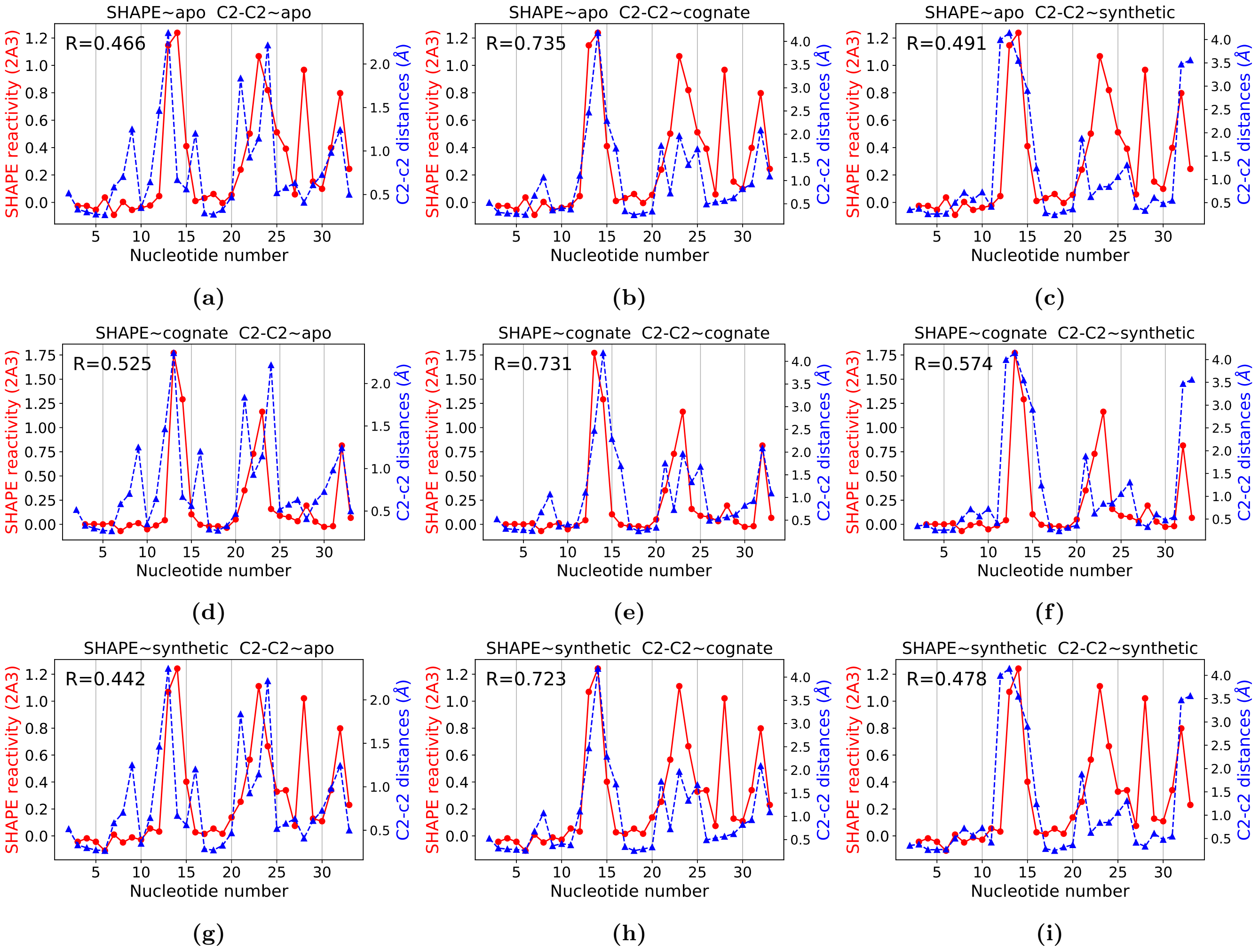


**Supplementary Figure 1:** SHAPE reactivities for the *Tt* $\text{PreQ}_1$ riboswitch aptamer obtained using the 2A3 reagent (red circles joined by solid lines) are compared with the fluctuations of C2 − C2 distances (blue triangles joined by dashed lines) for the $\text{PreQ}_1$ riboswitch. Respective columns correspond to fluctuations of C2 − C2 distances from different systems, while rows correspond to SHAPE-MaP measurements for different systems. Pearson correlation coefficients $R$ are shown in upper left corners.

### 9. AI-AUGMENTED ENHANCED SAMPLING METHOD RAVE

The unbinding of ligands from receptors such as proteins or nucleic acids is a prototypical example of a rare event in molecular simulations because its time scale is far beyond that can be reached by MD simulations. To study the unbinding process, here we combine a suite of advanced simulation approaches, namely metadynamics, RAVE, and AMINO[23–26] to enhance the sampling of rare events in an automated and robust manner.

In metadynamics, a time-dependent biasing potential as function of a pre-selected low dimensional order parameter (OP) is added to the underlying Hamiltonian of the system at a given frequency to enhance the fluctuation along this OP. As the external bias potential accumulates during the simulation, we expect the system will be able to cross different energy barriers and the trajectory to visit different metastable states. The net output of a typical metadynamics simulation is the free energy of the system along the biased variable or through a reweighting procedure along any other degree of freedom. In our study, we apply a variant of metadynamics called well-tempered metadynamics. In well-tempered metadynamics the height of the Gaussian kernel, instead of being a constant, decreases during the simulation whenever any point is revisited. This strategy has been shown to have better convergence property.[27]

While in principle metadynamics is guaranteed to yield the true free energy for any choice of biasing OP,[27] in practice the efficiency of metadynamics relies significantly on the choice of the OP being biased.[28] The efficiency is expected to be maximal if the true reaction coordinate (RC) for the process under investigation was chosen as the biasing OP. In this case all relevant fluctuations are enhanced through metadynamics, and any remaining modes are either fast or irrelevant; however, the choice of biasing OP is still an open question in the field. For instance, for a complex system consisting of a massive number of atoms, identifying the slow mode is computationally expensive. For example, even though the isocommittor is considered an ideal definition of slow mode, its calculation is prohibitively expensive for it to be practical as a biasing variable.[29] More generally, learning good biasing OPs requires access to trajectories that have already sampled the rare events of interest. In other words, effectively, one faces difficult a chicken-versus-egg problem, where learning good biasing variables needs access to sampling of the rare events of interest, but without access to good biasing variables such rare events of interest can simply not be sampled.

In this study, we use our recent methods that circumvent this coupled problem through the use of AI. Specifically, we make use of the past-future information bottleneck (PIB) framework to efficiently learn the useful but slowly varying features from a given high dimensional input dataset.[25,26,30] This PIB approximates the true RC, and is used as a biasing variable in next round of MD, through metadynamics or some other biasing framework, to generate new data from which a new, more accurate RC can be learnt. The iteration between generating data through MD and learning the RC through the PIB framework continues until further rounds of iteration do not improve the RC or the sampling. This combined protocol is named "Reweighted Autoencoded Variational Bayes for Enhanced sampling (RAVE)". The RC in RAVE is constructed as an interpretable, linear combination of a very large number of candidate OPs, which in the case of riboswitch-ligand dissociation comprise all nucleic acid-ligand contacts. As the number of all such contacts can be quite large, we use another pre-screening method "Automatic mutual information noise omission (AMINO)"[24] for pre-screening these candidate OPs, reducing the redundant ones that carry very similar information as each other through the use of a mutual information based metric. We provide further theoretical and implementation details of AMINO, RAVE and metadynamics.

We started with a dictionary of pre-selected OPs dictionary that contains the OPs relevant to our study. s However, we would like to note that we do not need to use all these OPs to describe the reaction process, as many of these OPs will likely be highly correlated and not capturing different physics. Therefore we used AMINO to pick out the most representative OPs from OP dictionary. AMINO embeds OPs from the dictionary into a space with an information theory based distance metric. More specifically OPs with high mutual information will be closer in the embedding space thus forming a cluster. Since OPs from the same cluster share similar information, only OPs in the center of each cluster are picked by AMINO to reduce redundancies.

To study ligand unbinding from riboswitch, we define the OP dictionary as all the pairwise distances between centers of riboses from individual nucleotides and heavy atoms from ligand. This amounts to a total of 429 and 627 trial OPs for the cognate and synthetic ligands respectively. For each system, we use four $500ns$ unbiased MD to calculate the distances. AMINO picks seven ligand-riboswitch distances for $\mathrm{PreQ}_1$ bound system and three ligand-riboswitch distances for synthetic ligand bound.

The OPs selected by AMINO are then fed into our iterative AI-MD scheme RAVE.[25,26] The central idea in RAVE is to learn a low dimensional representation, referred to as predictive information bottleneck (PIB), that is minimally complex but maximally predictive of the time evolution of the system, here characterized by the OPs. Identifying the most predictive combination of these OPs space will lead to the learning of features that persist with time. Enhancing fluctuations along such a RC will then lead to increased exploration of the system, which can then be fed back into the AI for learning another round of improved RC. Starting with the initial 2 $\mu s$ long unbiased trajectories, with every round of AI in RAVE here we learn a 1-dimensional RC approximation as a linear combination of these input OPs, which is then used as a biasing variable in the next round of metadynamics based biased simulation. As the MD-machine learning-MD iterations go on, the quality of the RC and ergodicity of sampling will keep improving. We finally stop this process when the quality of sampling does not improve with further training. This amounts four rounds of RAVE for the cognate and synthetic liganded systems.

### 10. NETWORK ARCHITECTURE AND HYPERPARAMETER SETTING FOR ENHANCED SAMPLING SIMULATION

#### 1. *AMINO*

We use AMINO[24] as a pre-processing method to reduce the number of order parameters fed to RAVE. AMINO achieves the goal of reducing the redundancy of the OP set by using the idea of clustering. AMINO uses a mutual information-base distance metric to quantify the distances between OPs, i.e., those OPs that share similar information will be closer to each other. Base on this distance metric, k-medoids clustering is use to group each OP into different clusters and the OP from the center of each group is selected as output to represent other OPs in the same group. The number of groups is determined by meausuring the rate–distortion functions.

To describe the relative position of ligands to the riboswitch, we choose the pairwise distance between each heavy atom from ligand and the center of mass of each ribose of individual nucleotide. Four 500ns unbiased trajectory is used as the input to learn the reduced set of OPs. The reduced set of OPs for riboswitch bound with cognate ligand and riboswitch bound with synthetic ligand are indicated in Fig. 3 in the main text.

#### 2. RAVE

In RAVE, a neural network with a linear encoder is trained to minimize the objective function

$$\mathcal{L}_{PIB} \equiv -I(\boldsymbol{\chi}; \mathbf{X_{\Delta t}}) + \alpha \mathbf{I}(\mathbf{X}; \boldsymbol{\chi}) \quad (1)$$

Here $\mathbf{X}$ is a vector of OPs that are used to describe the state of the system and $\mathbf{X_{\Delta t}}$ represent that state of system after time delay $\Delta t$. $\boldsymbol{\chi}$ is learned by the encoder as a linear combination of $\mathbf{X}$. $I(x, y)$ is the mutual information between two random variables $x$ and $y$. By minimizing the objective function, the information shared between $\boldsymbol{\chi}$ and $\mathbf{X}$ will be reduced while maintaining the information shared between $\boldsymbol{\chi}$ and $\mathbf{X_{\Delta t}}$. As a consequence, $\boldsymbol{\chi}$ will capture the slow mode of the system as it contains the features that are essential to describe dynamics of the system. We use the same network structure as in previous applications of RAVE.[26] We first set $\alpha = 0$ as we have a simple linear encoder which has already largely reduced the information shared between $\boldsymbol{\chi}$ and $\mathbf{X}$. The linear encoder was a densely connected layer without activation function or bias. A normalization constrain was also added to network parameter to reduce degeneracy as scaling the linear combination coefficients by any factor will not change the learned RC. The decoder consisted of two hidden layers with 128 nodes in each and an output layer. Exponential linear unit was used as the activation function for hidden layers and RMSprop algorithm[31] was used to optimize the network parameters.

#### 3. Metadynamics

In the well-tempered Metadynamics, the initial bias high was set to be 1 kJ/mol. The variance of Gaussian potential was set to be half of the standard deviation of the RC in the unbiased MD simulation. The bias was deposited every 4 ps, and the bias factor was set to be 10 to control the decrease of bias high during the simulations. In each round of the iteration, we ran 8 simulations with randomly initialized velocities. The simulations stopped when the coordination number was lower than a threshold, which indicated the dissociation of ligands from the riboswitch. These trajectories were mixed together to train the RC for the next round of metadynamics simulation. Before the final round, it took around 200 ns for ligands to dissociate with the suboptimal RCs. In the final production runs, dissociation of ligands usually took less than 50 ns.

### 11. PREDICTING CRITICAL NUCLEOTIDES FOR MUTAGENESIS EXPERIMENTS

With the biased MD, we can have more ergodic sampling of possible interaction poses of these systems and make predictions of mutation that can bring different binding affinity changes. Our guiding principle here is that the nucleotides which exhibit different behavior when a given ligand unbinds, should play differing roles in determining the binding affinity, as their interactions with the ligand should play a critical role in determining the bound state ensemble. We can directly check the RMSD of each reach nucleotide as the two ligands unbind during the biased MD simulation.

For each nucleotide $i$ for a given ligand, we can calculate its average RMSD given a specific value of coordination number $C$.

$$\mathrm{RMSD}_i(C) = \int \mathrm{RMSD} \cdot P_i(\mathrm{RMSD}|C) d\mathrm{RMSD} \quad (2)$$

Where $P_i(\mathrm{RMSD}|C)$ is the distribution of root mean square deviation of nucleotides $i$ given coordination number $C$. This distribution can be calculated from biased MD after appropriate reweighting.[27,28]

To emphasize movement of nucleotides induced by ligand dissociation, we subtract from the various RMSDs their corresponding mean values in the unliganded apo system from the long unbiased MD. The average RMSD of individual nucleotides as function of coordinate number are shown in Supplementary Fig. 2. Here we can ignore C1 and G33 because they lie in the flexible ends of the riboswitch. Then we categorize the rest of nucleotides into three cases. In the first case, the RMSD doesn't change much as both ligands disassociate, such as G5 and U6 which indicates that those residues do not undergo conformational change when either ligand unbinds. In the second case, for both the ligands the RMSD undergoes significant change when the coordination number decreases, such as N13 and N14, which indicates that those residues need to adopt new configurations when ligand leave the binding pocket. In the third case, the RMSD shows significant change during ligand dissociation only for one of the ligands. These are U22 and A32 which are highlighted by thicker lines in Supplementary Fig. 2. This case is the most interesting to us because these different structural changes indicate that those nucleotides interact with different ligands differently when they disassociate. We can also observed that for the cognate ligand, the dominant dissociation pathway was closer to U22, while for the synthetic ligand, it is closer to A32. Such a preference on unbinding pathways might be related to the more significant change in the RMSD profile for one of the ligands. I selected these two nucleotides for mutagenesis experiments and expected that those mutations would bring more significant changes to the binding affinity of one of the ligands while bringing a relatively more minor binding affinity change to another one.

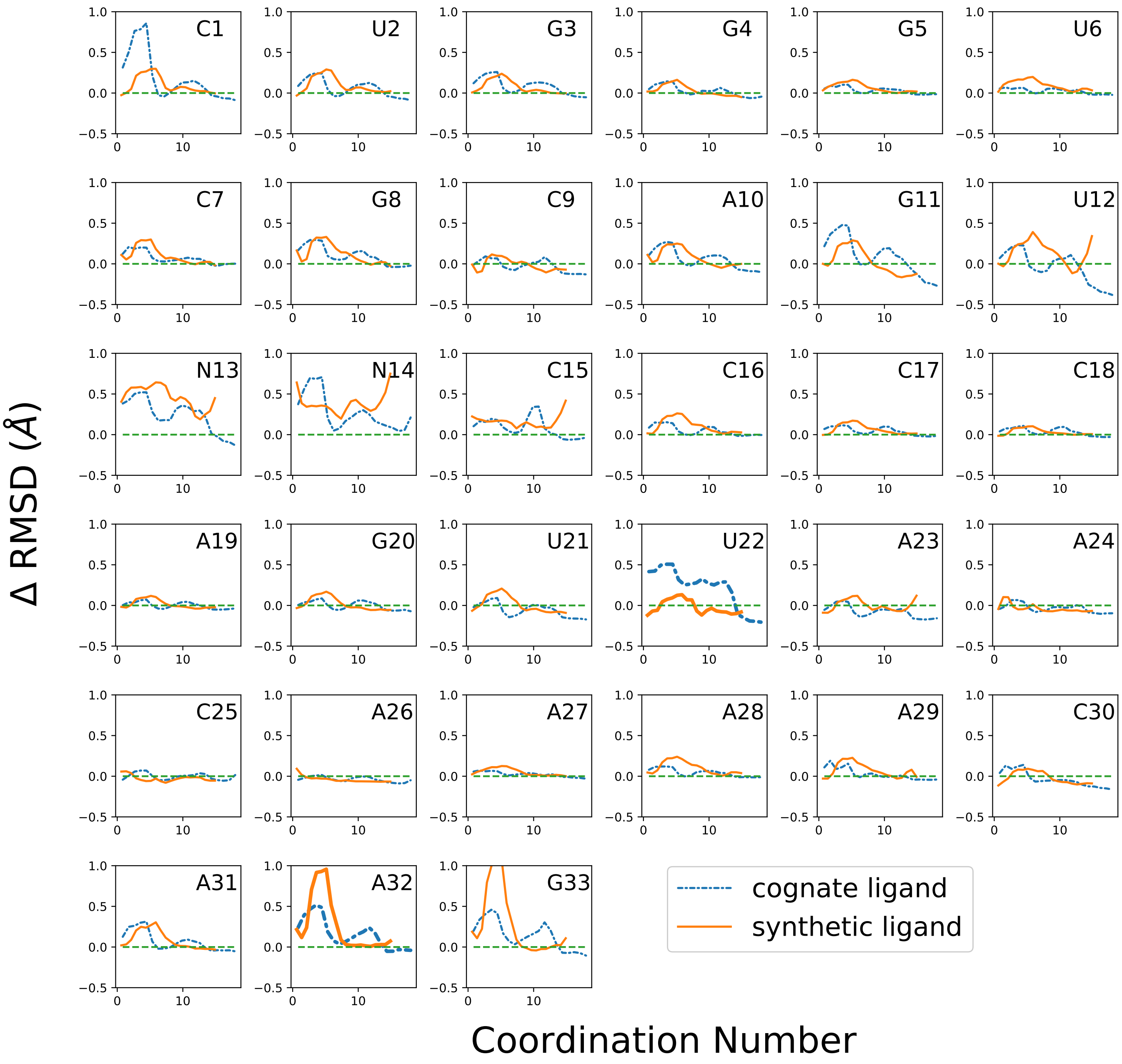


**Supplementary Figure 2:** The change of RMSD of each nucleotide during the dissociation process. RMSD as a functions of coordination number are shown by blue dashed line and orange solid line for systems with cognate ligand and synthetic ligand separately.

### 12. *IN SILICO* MUTAGENESIS EXPERIMENTS

We used PyMOL[32] to generate the mutants U22A and A32U for riboswitch with cognate or synthetic ligands. We performed four independent unbiased simulations for each system, with each 500ns long. The structures were saved every 10 ps. In the 500 ns simulations, the ligands stay in the bound pose. The statics of samples from these simulations can be used to illustrate the influence of mutations on the bound state ensemble. Supplementary Fig.3 shows the free energy profiles of these systems and clearly indicates the changes in the bound state ensemble brought about the mutations U22A and A32U.

To further trace the possible interactions that bring changes to the free energy profiles, we studied the hydrogen bond occupancy between ligand and nucleotides from riboswitch in $2\mu s$ long unbiased MD simulation. This captures the bound state ensemble better than just the crystal structure and shows how the mutations, even though distal, disrupt the hydrogen binding patterns. The number of hydrogen bonds is calculated by VMD[33] with an angle cutoff of 30 degrees and a distance cutoff $3.5\mathring{A}$. The hydrogen bond occupancy is defined as $\bar{\rho} = \sum_{i=1}^{n} N_i/n$ where $n$ is the number of total frame and $N_i$ is the number of hydrogen bonds in frame $i$. Supplementary Fig.4 shows that different hydrogen bonds between ligand-nucleotide pairs are formed in the bound state ensemble before and after the mutations. For example, the hydrogen bonds between cognate ligand and C16 disappeared after the mutation and the hydrogen bonds between cognate ligand and A28 mainly showed up in the A32U mutant. Also, for synthetic ligand, the hydrogen occupancy with U6 largely increases after the mutations while that with G4 decreases.

Here we would like to note that these *in silico* mutagenesis experiments only serve the purpose of validating the effect of muation on the bound state ensemble but not trying to give a thorough study of how the mutations change the bound state ensemble. This is because the $2\mu s$ unbiased simulation will not provide an ergodic sampling even just for the bound states ensemble. Also, besides hydrogen bond, there are other interactions such as $\pi$-$\pi$ stacking which are also important in terms of understanding ligand-riboswitch interactions but we are not going to discuss the effect of mutations on those interactions in this work. Thus these hydrogen bonding and free energy analyses give a sense of why these mutations might matter. It is however only through our change in RMSD based and dissociation pathway based analyses, as reported in main text, that we can pinpoint the exact variation in behavior expected for the cognate and synthetic ligand systems with these two mutations.

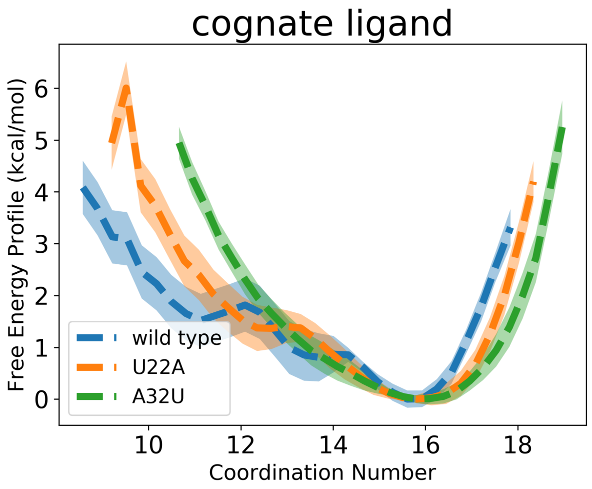


**(a)**

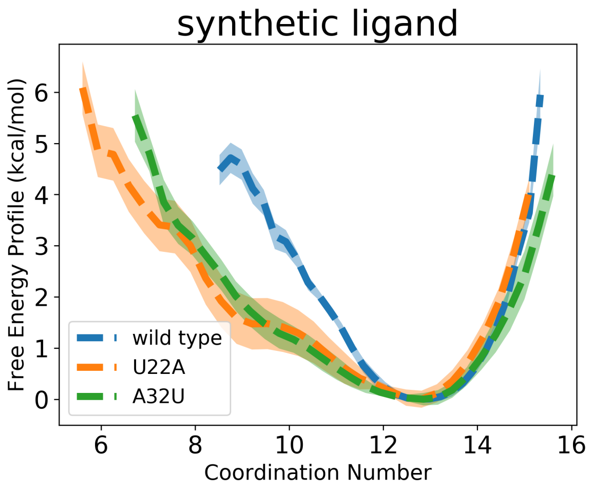


**(b)**

**Supplementary Figure 3:** Free energy profile from $2\mu s$ unbiased simulation for wild tpye and mutated systems with (a) cognate ligand and (b) synthetic ligand. The error bars are indicated by the shaded region around each curve.

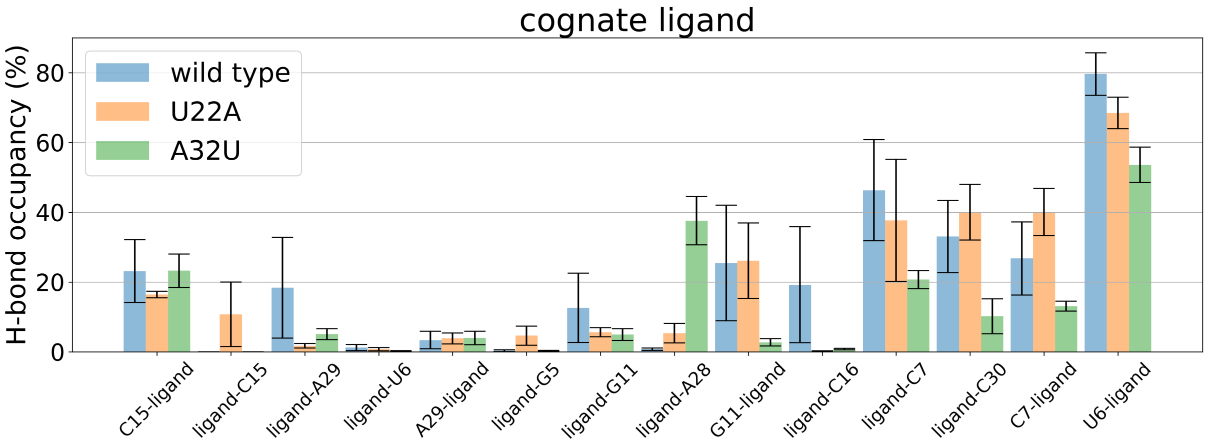


**(a)**

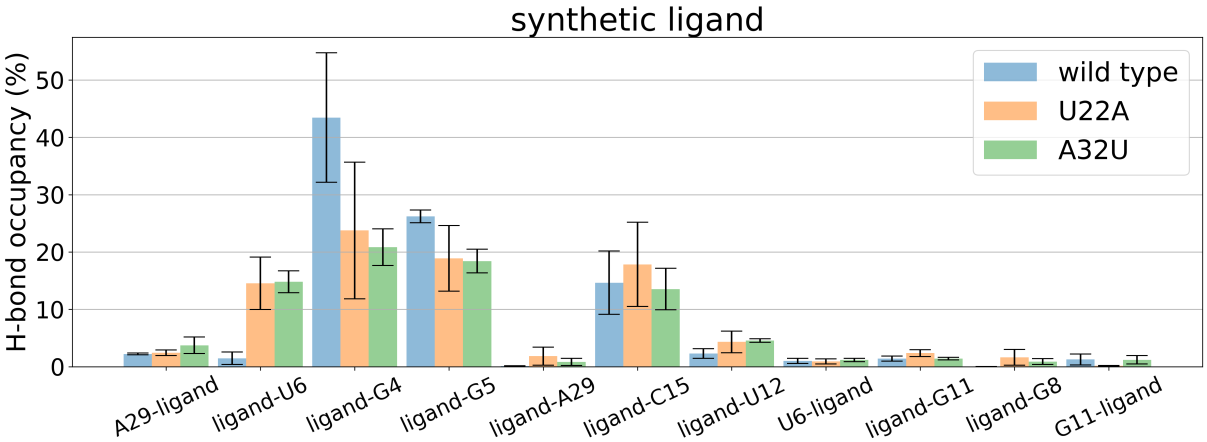


**(b)**

**Supplementary Figure 4:** Hydrogen bond occupancy for wild type and mutated systems calculated through $2\mu s$ long unbiased MD simulation with (a) cognate ligand and (b) synthetic ligand. The labels on the x-axis define the hydrogen bond pairs with the format "donor-acceptor".
